# Elimination of senescent cells by mechanical cell competition

**DOI:** 10.64898/2026.05.18.725837

**Authors:** Yuping Pan, Mattheus Xing Rong Foo, Vinod S/O Prabhakaran, Kashish Jain, Pakorn Kanchanawong, Oliver Dreesen, Yusuke Toyama

## Abstract

Cellular senescence, a hallmark of aging, leads to the accumulation of apoptosis-resistant cells that compromise tissue homeostasis. While senescent cells are known to influence neighboring cells through the senescence-associated secretory phenotype (SASP), the precise nature of the interactions between senescent and normal cells remains elusive. Here we show that progerin-induced senescent cells undergo apoptosis when co-cultured with normal cells. This elimination requires direct cell-cell contact and is mediated by the JNK and p38-MAPK pathways, leading to p53 upregulation and p21 downregulation in progerin-expressing cells. Furthermore, neighboring normal cells exert persistent mechanical compression on progerin-expressing cells prior to their elimination, consistent with mechanical cell competition. In contrast, p16-induced senescent cells resist elimination under the same co-culture conditions, maintaining high p21 levels. Our findings reveal a non-cell-autonomous mechanism for senescent cell clearance, providing new insights into the maintenance of tissue homeostasis during aging.

## Introduction

Aging and various stressors, including DNA damage, oncogene activation, and oxidative stress, lead to the induction of cellular senescence in tissues. Cellular senescence is a hallmark of aging and plays a central role in its progression^1^. Senescent cells are characterized by irreversible cell cycle arrest and resistance to apoptosis, allowing them to persist in tissues over time^2^. Cellular senescence is primarily regulated by two pathways: the p53/p21 pathway and the p16^INK4A^ pathway^3^. Overexpression of progerin, a mutant form of lamin A associated with premature aging disorders^4^, can also induce cellular senescence through the DNA damage response and p53 activation^5^. A major feature of senescent cells is the senescence-associated secretory phenotype (SASP), a complex mix of pro-inflammatory cytokines, chemokines, proteases, and growth factors^6^. SASP factors can induce senescence in neighboring normal cells, amplifying the detrimental impact of senescent cell accumulation^7^.

At the early stages of aging, not all cells undergo cellular senescence simultaneously. Rather, senescent cells arise sporadically among normal cells, creating a heterogeneous tissue environment where senescent and non-senescent cells coexist. As senescent cells accumulate over time, this heterogeneity increases, disrupting the balance of cell populations and compromising tissue homeostasis^6^. Tissues are known to employ homeostatic mechanisms to cope with cellular heterogeneity^8^. One such mechanism is cell competition, a process by which cells of different fitness levels interact, leading to the elimination of less-fit cells by their neighbors^9^. Cell competition has been extensively studied in *Drosophila*^10^ and is increasingly recognized in mammalian tissues, particularly in the context of oncogenic transformation^11–14^.

Cell competition can be mediated through juxtacrine signalling via direct cell-cell contact^15,16^, secreted signalling through paracrine factors^17,18^, and mechanical signalling through forces exerted by neighboring cells^19,20^. Among these, recent studies have demonstrated that mechanical compression by normal cells can eliminate unfit cells, a process termed mechanical cell competition^19,20^. However, whether such competitive interactions operate between senescent and normal cells, and how these interactions influence the fate of senescent cells, remain poorly understood.

Here, using an *in vitro* co-culture system of progerin-induced senescent cells and normal cells, we found that senescent cells undergo apoptosis through a contact-dependent mechanism involving JNK and p38-MAPK pathways and mechanical compression by neighboring normal cells, consistent with mechanical cell competition.

## Results

### Progerin-induced senescent cells undergo apoptosis when co-cultured with normal cells

To investigate the interaction between senescent and normal cells, we established Madin-Darby Canine Kidney (MDCK) epithelial cells overexpressing progerin using a doxycycline-inducible system, with mCherry expressed independently of progerin as a marker for overexpressing cells (pLVX-TetOne-mCherry-Puro-Progerin, hereafter referred to as mCherry-Progerin, Methods, Fig. S1A). Upon doxycycline treatment after cells formed a confluent tissue, progerin expression and hallmarks of cellular senescence, including increased cell size, elevated SA-β-gal, increased γH2A.X, and reduction of Lamin B1 and Ki67, were apparent (Fig. S1B-H). To examine the fate of progerin-induced senescent cells next to normal cells, we co-cultured normal and mCherry-Progerin cells at a ratio of 50:1 on an elastic polydimethylsiloxane (PDMS) gel substrate (∼15kPa) coated with fibronectin (Methods). Once a confluent monolayer was formed, cells were treated with doxycycline (Fig. S1I) and subjected to time-lapse imaging (Methods).

To our surprise, progerin-induced senescent cells in co-culture were eliminated and displayed hallmark features of apoptosis^21^, including cellular shrinkage, plasma membrane blebbing, the formation of apoptotic bodies, and cell extrusion (Fig. 1A, Supp. Video 1). To confirm the modality of observed cell elimination, we examined the activity of cleaved caspase-3, a key effector enzyme in the apoptotic pathway^21^. Cleaved caspase-3 was detected in mCherry-Progerin cells while they were still part of the tissue (Fig. 1B), indicating that the elimination process is apoptosis rather than anoikis or live cell extrusion. We quantified the fraction of apoptotic cells (Methods) and found that progerin-induced senescent cells in co-culture underwent significantly more apoptosis compared to the control without doxycycline treatment (Fig. 1C-D). This observation is counterintuitive, as senescent cells are typically resistant to apoptosis. To further validate this finding, we employed two independent strategies. First, we established MDCK cells expressing mNeonGreen-tagged progerin in a doxycycline-inducible manner (mNG-T2A-Progerin, hereafter referred to as mNG-Progerin, Methods, Fig. S2A). Three days of doxycycline treatment led to progerin expression (Fig. S2B) and cellular senescence, evidenced by multiple senescence characteristics (Fig. S2C-E). Similar to mCherry-Progerin cells, mNG-Progerin cells in co-culture underwent apoptosis (Fig. 1D-E, Supp. Video 2). Second, to confirm that this phenomenon is not restricted to MDCK cells, we established HK-2 human kidney cells expressing mNG-T2A-Progerin (hereafter referred to as HK-2 mNG-Progerin, Methods). Consistent with our other observations, HK-2 mNG-Progerin cells were eliminated under co-culture conditions (Fig. S2F). Together, with the confirmation that the level of progerin expression is within physiological range^22^ (Fig. S2G), and consistent observations across two different constructs and two cell types, we concluded that progerin-induced senescent cells undergo apoptosis when they are surrounded by normal cells.

**Fig. 1.**
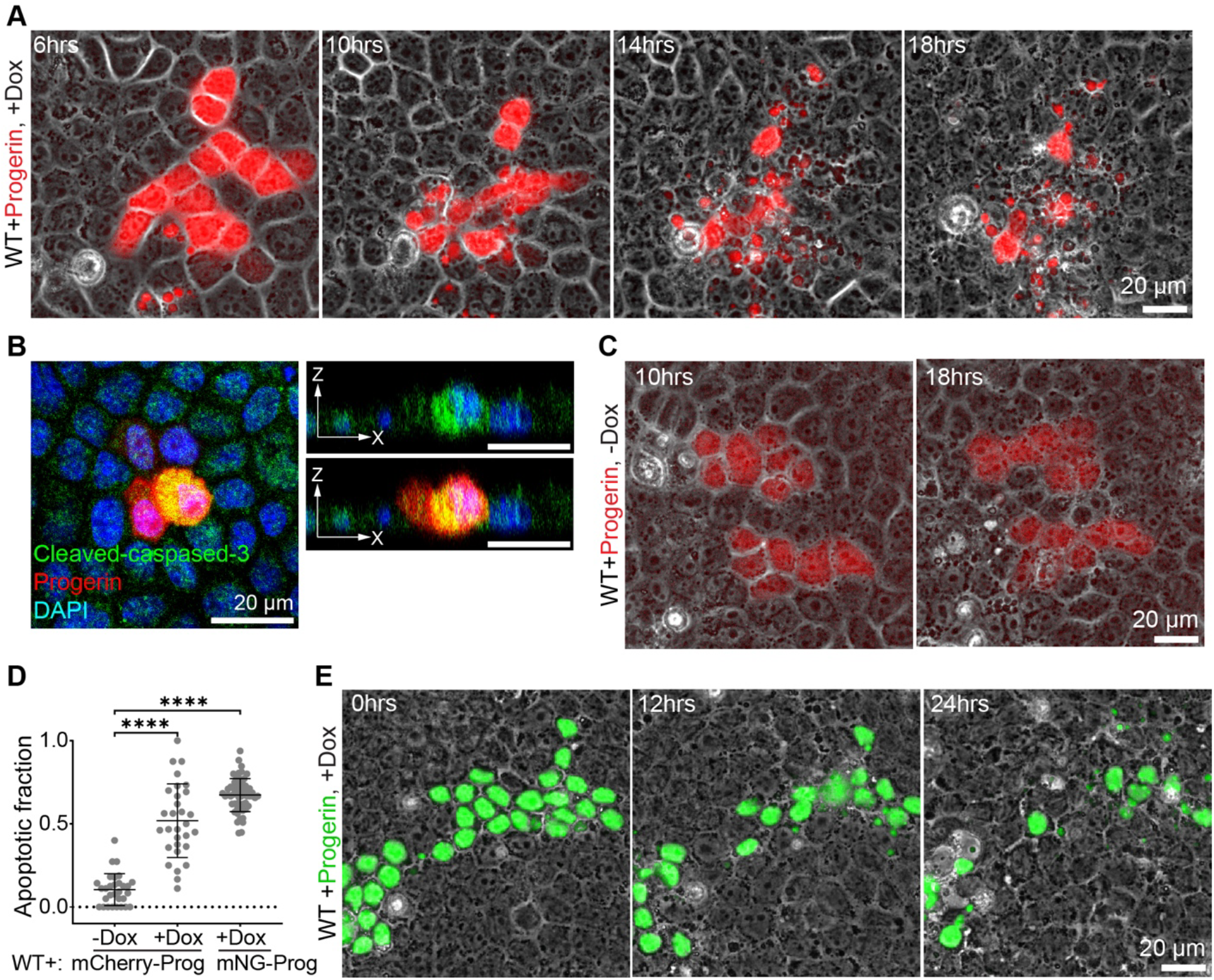
Progerin-induced senescent cells undergo apoptosis in co-culture with normal cells. (A) Time-lapse images of mCherry-Progerin cells (red) co-cultured with normal cells after doxycycline treatment. (B) Immunofluorescence images of cleaved caspase-3 (green) in co-culture tissue with doxycycline treatment. Nuclei were stained with Hoechst 33342 (blue). Cross-sectional images show cleaved caspase-3-positive mCherry-Progerin cells maintained within the tissue. (C) Time-lapse images of mCherry-Progerin cells (red) co-cultured with normal cells without doxycycline treatment. (D) Apoptotic fraction of mCherry-Progerin cells with and without doxycycline treatment, and mNG-Progerin cells with doxycycline treatment, in co-culture with normal cells. (E) Time-lapse images of mNG-Progerin cells (green) co-cultured with normal cells after doxycycline treatment. Data are presented as mean ± SD, (independent repeats, n = 3, N=30, 30, and 60). Kruskal-Walli’s test and Dunn’s multiple comparisons test are used. ^**^*p*<0.0001

### Cell-cell contact with normal cells is required for apoptosis in progerin-induced senescent cells

To investigate whether physical contact with normal cells is required for the observed elimination of progerin-induced senescent cells, we first compared the fraction of apoptotic cells when mCherry-Progerin cells alone formed a monolayer (Fig. 2A, Supp. Video 3). The fraction of apoptotic cells in the mCherry-Progerin alone condition was as low as the control (i.e., no doxycycline treatment), and significantly lower than the co-culture condition (Fig. 2B). To perturb cell-cell contact between progerin-induced senescent cells and normal cells, we co-cultured mCherry-Progerin cells with E-cadherin knockout (KO) cells^23^. We found a reduction in the elimination of progerin-expressing cells compared to the condition where normal cells were the neighbors (Fig. 2C, Supp. Video 4), further supporting the idea that cell adhesion is crucial for the elimination of progerin-induced senescent cells. To investigate whether long-range signalling, including secretory factors, plays a role in this elimination process^24^, we cultured senescent and normal tissues side by side using a culture-insert two-well system (Methods, Fig. 2E). We first allowed the two cell populations to establish cell-cell contacts, then administered doxycycline for three days. Time-lapse imaging was performed for two days, and the fraction of apoptotic cells was measured at two locations: progerin cells at the tissue interface, and progerin cells far from the interface (Fig. 2E). The fraction of apoptotic cells was higher at the interface compared to the interior of the tissue, which was comparable to the fraction in progerin-expressing cells cultured alone (Fig. 2E-F). Together, our data demonstrated that long-range secreted factors from normal cells are unlikely to contribute to the elimination, but that cell-cell contact is required for the induction of apoptosis in progerin-expressing cells under co-culture conditions.

**Fig. 2.**
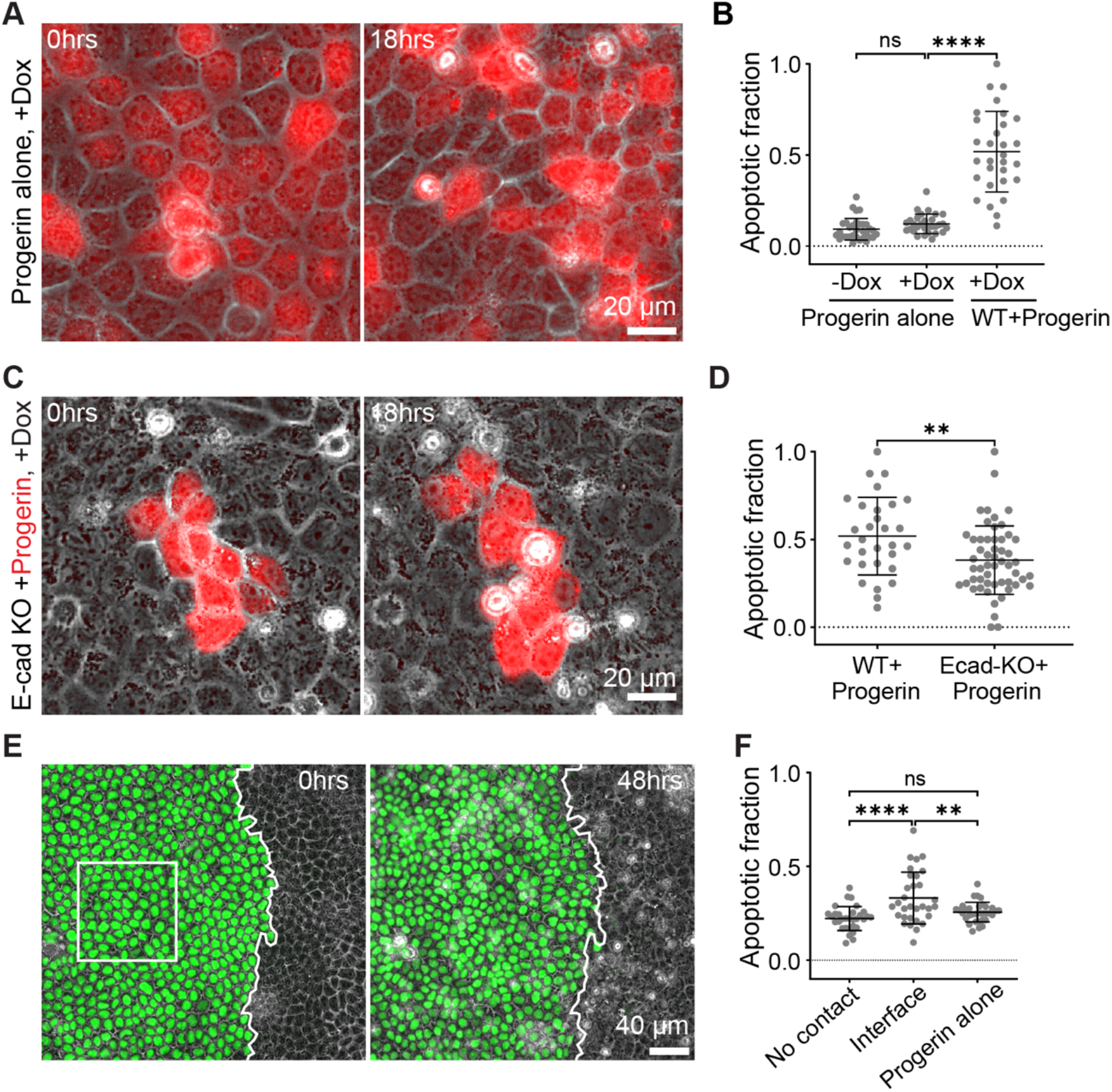
Cell-cell contact with normal cells is required for apoptosis in progerin-induced senescent cells. (A) Time-lapse images of mCherry-Progerin cells cultured alone with doxycycline treatment. (B) Apoptotic fraction of progerin cells cultured alone with and without doxycycline treatment, and in co-culture with normal cells with doxycycline treatment (shown in Figure1D). (C) Time-lapse images of mCherry-Progerin cells (red) co-cultured with E-cadherin knockout (KO) cells after doxycycline treatment. (D) Apoptotic fraction of progerin cells co-cultured with E-cadherin KO cells compared to co-culture with normal cells (shown in Figure1D). (E) Time-lapse images of a culture-insert two-well system at 0 and 48 hours. mNG-Progerin cells at the interface with normal cells are indicated. (F) Apoptotic fraction of mNG-Progerin cells at the interface with normal cells, in regions without contact with normal cells, and in mNG-Progerin cells cultured alone. Data are presented as mean ± SD. Kruskal-Walli’s test and Dunn’s multiple comparisons test are used in (B), (independent repeats, n = 3, N = 30, 30, and 30). Unpaired t test, two-tailed is use in (D), (independent repeats, n = 3, N = 30, 54). Kruskal-Walli’s test, and Dunn’s multiple comparisons test are used in (B), (independent repeats, n = 3, N = 30, 30, 30). One-way ANOVA multiple comparison was used in (F), (independent repeats, n=4 for culture insert and n = 3 for progerin alone, N = 31, 31, and 33). ns, not significant; **, p < 0.01; ****, p < 0.0001.

### Apoptosis in progerin-induced senescent cells is JNK and p38-MAPK dependent

To understand the mechanisms underlying apoptosis in progerin-induced senescent cells under co-culture conditions, we performed a small-molecule inhibitor screen (Fig. 3A, Supp. Table 1). Co-cultured tissues of mCherry-Progerin and normal cells (50:1) were treated with doxycycline along with each inhibitor, followed by time-lapse imaging for 24 hours (Fig. 3A). Inhibitors targeting the mitogen-activated protein kinase (MAPK) pathways, including c-Jun N-terminal kinase (JNK) (SP610025), p38-MAPK (PD169316), and extracellular signal-regulated kinase (ERK) (U0126), along with Tumour Necrosis Factor-α (TNFα) inhibitor and Cdc42 inhibitor (ML-141), showed inhibition of apoptosis compared to the DMSO control (Fig. 3B). Among MAPKs, ERK is known for primarily antiapoptotic effects^25^, and we therefore focused on JNK and p38-MAPK, which are known to promote apoptosis among other cellular responses^26^. TNFα and Cdc42 are also known to function upstream of JNK and p38-MAPK pathways^26^. To further confirm the involvement of JNK and p38-MAPK, we performed immunostaining of JNK (JNK1,2,3) and p38-MAPK (phosphorylated p38-MAPK) in co-culture tissue with inhibitor treatment. In mCherry-Progerin cells, both JNK and p38-MAPK were elevated in the nucleus compared to neighboring normal cells (Fig. 3B, C, F, H). The levels of JNK and p38-MAPK were also higher than those in mCherry-Progerin cells cultured alone (Fig. 3D-E, G-H). The levels of JNK and p38-MAPK showed no difference between co-culture and mCherry-Progerin alone tissues without doxycycline treatment (Fig. S3A-F). Collectively, our data indicate that JNK and p38-MAPK pathways are associated with apoptosis of progerin-induced senescent cells in co-culture tissue.

**Fig. 3.**
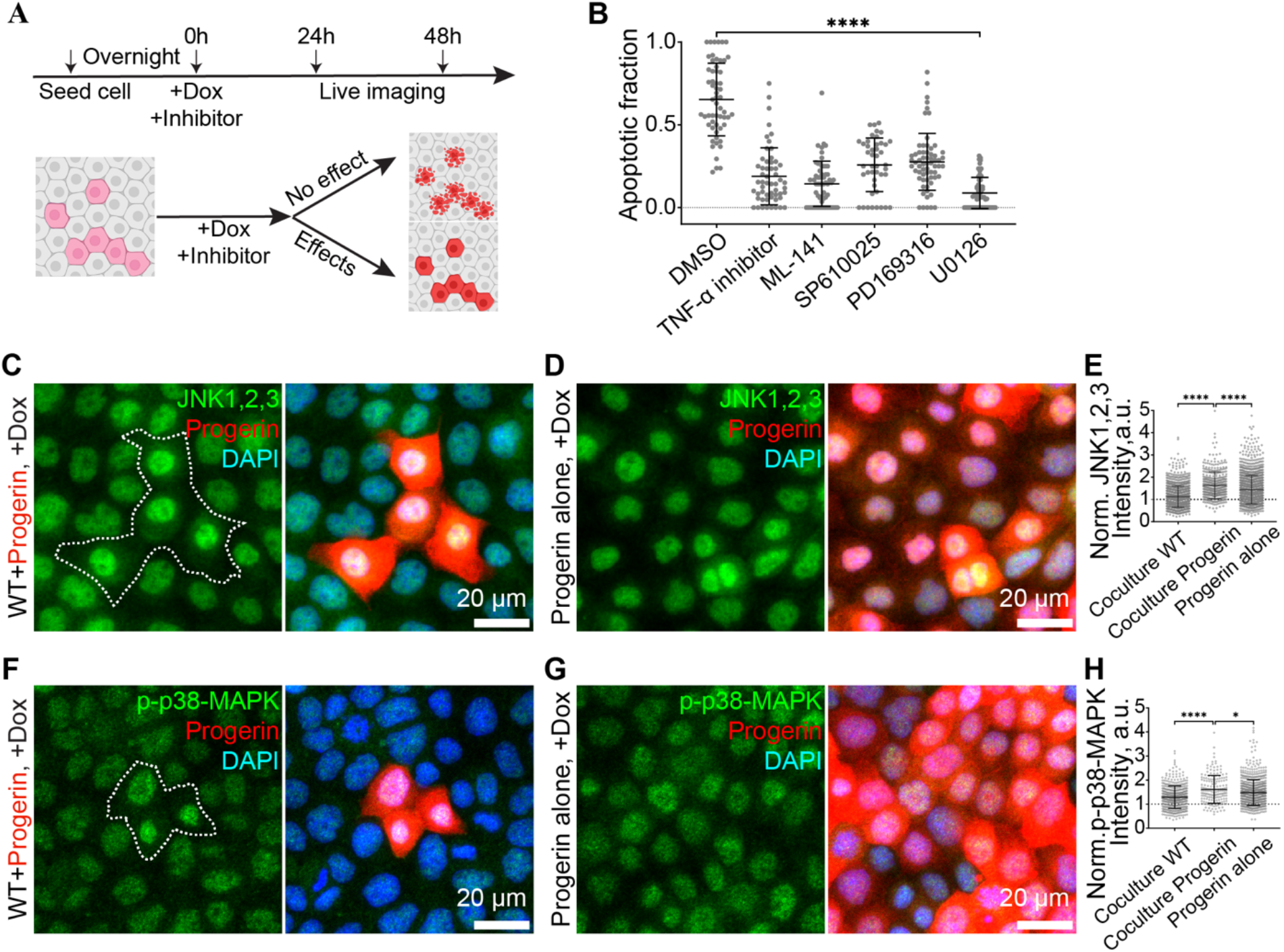
JNK and p38-MAPK are elevated in progerin-expressing cells in co-culture tissue. (A) Schematic of the small-molecule inhibitor screen. Co-cultured tissues were treated with doxycycline and each inhibitor, followed by time-lapse imaging. (B) Apoptotic fraction of progerin cells after treatment with DMSO, TNFα inhibitor, Cdc42 inhibitor (ML-141), JNK inhibitor (SP610025), p38-MAPK inhibitor (PD169316), and ERK inhibitor (U0126). (C) Representative immunofluorescence images of JNK1,2,3 (green) in co-culture tissue with doxycycline treatment. Progerin cells are indicated by dashed outlines. Nuclei were stained with Hoechst 33342 (blue). (D) Representative immunofluorescence images of JNK1,2,3 in progerin cells cultured alone with doxycycline treatment. (E) Quantification of normalized JNK1,2,3 fluorescence intensity in neighboring normal cells and progerin cells in co-culture, and progerin cells cultured alone. (F) Representative immunofluorescence images of p-p38-MAPK (green) in co-culture tissue with doxycycline treatment. (G) Representative immunofluorescence images of p-p38-MAPK in progerin cells cultured alone with doxycycline treatment. (H) Quantification of normalized p-p38-MAPK fluorescence intensity in neighboring normal cells and progerin cells in co-culture, and progerin cells cultured alone. Data are presented as mean ± SD. Kruskal-Walli’s test and Dunn’s multiple comparisons test are used in (B), (independent repeats, n = 3 and (N = 4 for DMSO and ML-141), N= 58, 55, 60, 44, 64, 60). Kruskal-Walli’s test and Dunn’s multiple comparisons test are used in (E), (independent repeats, n = 4, N = 1109, 411, and 1555). Kruskal-Walli’s test and Dunn’s multiple comparisons test was used in (H), (independent repeats, n=3, N = 472, 175, and 742). ^*^*p*<0.05, ^**^*p*<0.0001.

### p53 upregulation and p21 downregulation are associated with apoptosis of progerin-induced senescent cells

Our data so far indicate that JNK and p38-MAPK pathways are involved in the observed apoptosis of progerin-expressing cells. However, the downstream cellular responses of JNK and p38-MAPK pathways are wide-ranging, including differentiation and proliferation ^27,28^, and are not restricted to apoptosis. To confirm that the downstream consequence of elevated JNK and p38-MAPK in the nucleus is indeed apoptosis, we further examined the downstream effectors of JNK and p38-MAPK pathways. We first examined the level of the pro-apoptotic factor p53 by immunostaining. Progerin-expressing cells in the co-culture condition showed increased p53 compared to neighboring normal cells and to progerin-expressing cells cultured alone (Fig. 4A-C). We next examined p21, a downstream effector of p53 known for regulating the cell cycle, promoting senescence, and inhibiting apoptosis^29^. p21 showed a decrease in progerin-expressing cells compared to neighboring normal cells and to progerin-expressing cells cultured alone (Fig. 4D-F). The levels of p53 and p21 without doxycycline treatment were similar to the control condition (Fig. S4A-F). To further investigate whether these changes in p21 are downstream of the JNK and p38-MAPK pathways, co-culture tissues were treated with the JNK and p38 inhibitors SP610025 and PD169316, respectively, and the level of p21 was monitored. We found that although p21 in progerin-expressing cells was still lower than in neighboring normal cells (Fig. 5A-D), the degree of reduction was mitigated in inhibitor-treated conditions compared to untreated co-culture tissue (Fig. 5E). These results suggest that JNK and p38-MAPK pathways, in part, contribute to the reduction of p21 levels in co-culture conditions. Taken together, our data demonstrated that elevated JNK and p-p38-MAPK levels in progerin-expressing cells are associated with increased p53 and reduced p21 levels in co-culture tissues, thereby promoting apoptosis in progerin-induced senescent cells.

**Fig. 4.**
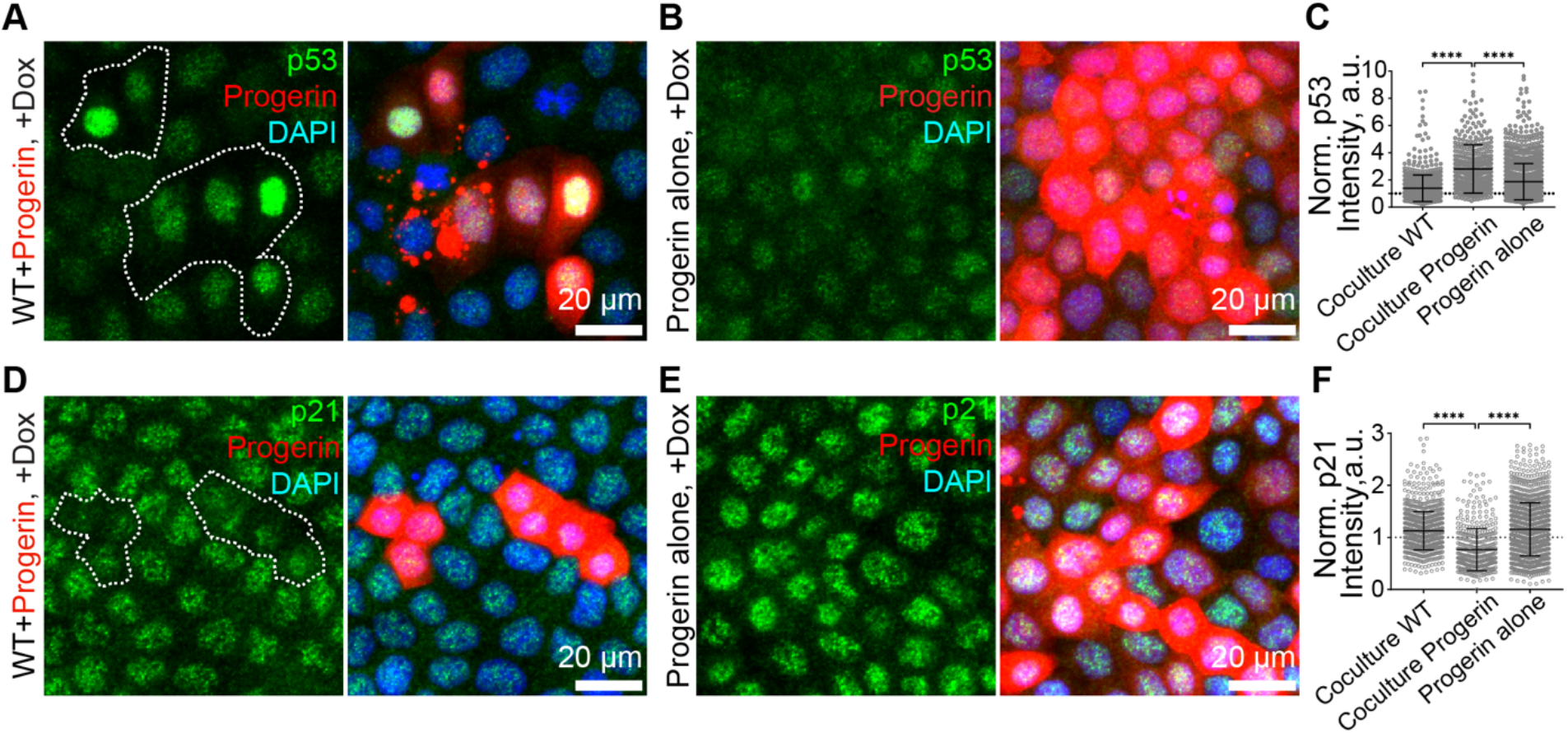
p53 increases and p21 decreases in progerin-expressing cells in co-culture tissue. (A) Representative immunofluorescence images of p53 (green) in co-culture tissue with doxycycline treatment. Progerin cells are indicated by dashed outlines. Nuclei were stained with Hoechst 33342 (blue). (B) Representative immunofluorescence images of p53 in progerin cells cultured alone with doxycycline treatment. (C) Quantification of normalized p53 fluorescence intensity in neighboring normal cells and progerin cells in co-culture, and progerin cells cultured alone. (D) Representative immunofluorescence images of p21 (green) in co-culture tissue with doxycycline treatment. (E) Representative immunofluorescence images of p21 in progerin cells cultured alone with doxycycline treatment. (F) Quantification of normalized p21 fluorescence intensity in neighboring normal cells and progerin cells in co-culture, and progerin cells cultured alone. Data are presented as mean ± SD. Kruskal-Walli’s test and Dunn’s multiple comparisons test are used in (C), (independent repeats, n = 4, N = 1314, 440, and 1421). Kruskal-Walli’s test and Dunn’s multiple comparisons test was used in (F), (independent repeats, n = 4, N = 1040, 419, and 1569). ^**^*p*<0.0001

**Fig. 5.**
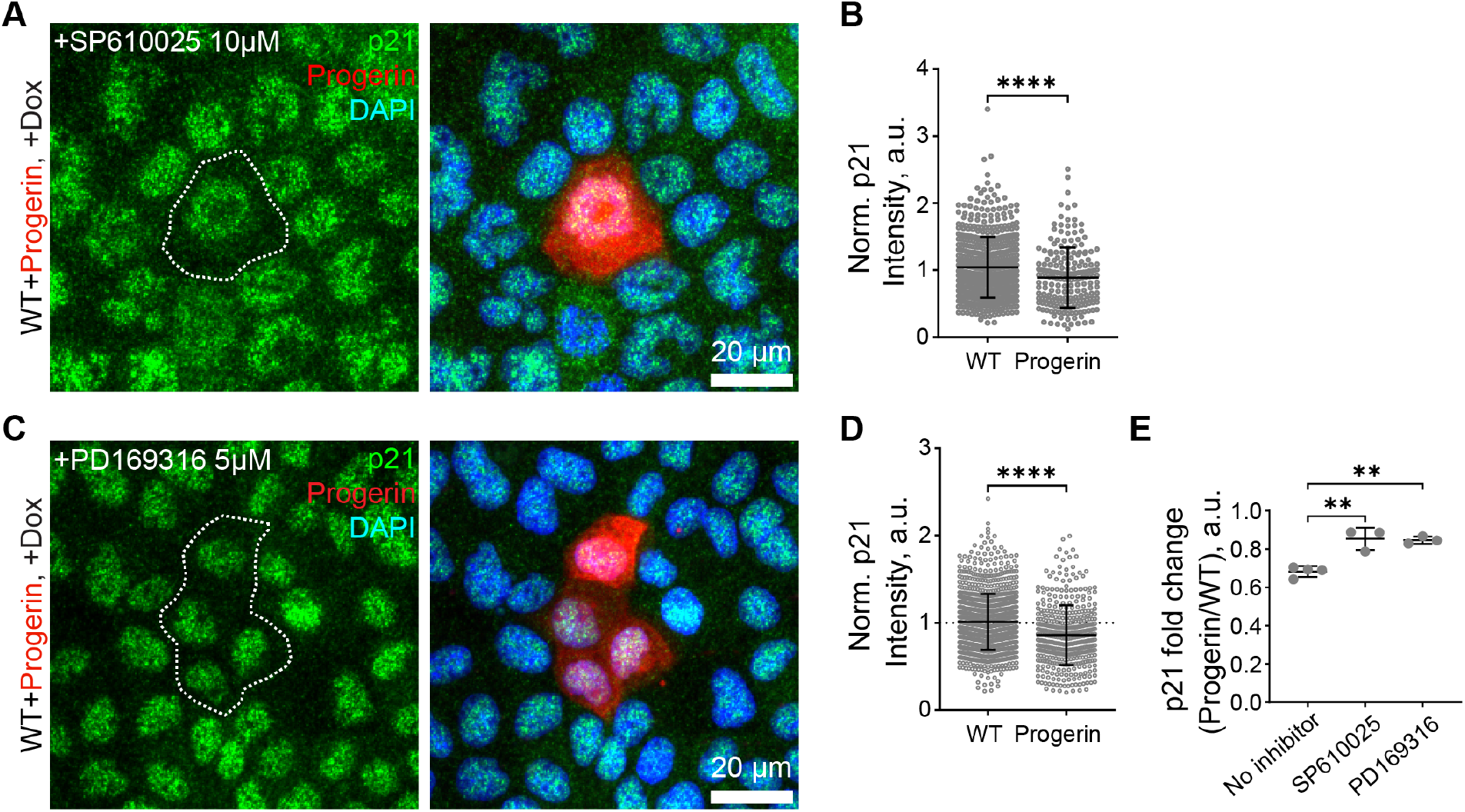
JNK and p38 inhibitors partially rescue p21 reduction in progerin-expressing cells. (A) Representative immunofluorescence images of p21 (green) in co-culture tissue treated with JNK inhibitor SP610025 and doxycycline. Progerin cells are indicated by dashed outlines. Nuclei were stained with Hoechst 33342 (blue). (B) Quantification of normalized p21 fluorescence intensity in neighboring normal cells and progerin cells in co-culture treated with SP610025. (C) Representative immunofluorescence images of p21 in co-culture tissue treated with p38-MAPK inhibitor PD169316 and doxycycline. (D) Quantification of normalized p21 fluorescence intensity in neighboring normal cells and progerin cells in co-culture treated with PD169316. (E) Fold change of p21 intensity (progerin/normal) in untreated, SP610025-treated, and PD169316-treated co-culture tissues. Data are presented as mean ± SD. The Mann-Whitney test is used in (B), (independent repeats, n = 3, N = 565, and 200). The Mann-Whitney test is used in (D), (independent repeats, n = 3, N = 1090, and 419). Ordinary one-way ANOVA multiple comparison was used in (E). ^*^*p*<0.05 ^**^*p*<0.0001.

### Mechanical compression by neighboring cells precedes apoptosis of progerin-expressing cells

Having established that JNK and p38-MAPK pathways are involved in the apoptosis of progerin-expressing cells, we next sought to identify what activates these pathways. We considered whether mechanical cell competition could underlie the observed elimination. It has been demonstrated that neighboring normal cells compress cells silenced for the polarity gene scribble (scrib^KD^), leading to activation of p38 and in turn elevation of p53, causing cell death^20^, similar to the progerin-induced cell elimination observed here. To test this hypothesis, we used traction force microscopy. Tissues were cultured on a PDMS gel substrate embedded with fluorescent beads, and traction forces exerted by the cells were calculated based on bead movement (Methods, Fig. 6A, Supp. Video 5) before and after the death of progerin-expressing cells. To evaluate whether progerin-expressing cells were compressed by neighboring normal cells, we analyzed the radial component of the traction force with respect to the progerin cell. The radial traction force pointing away from the progerin cell (positive radial traction force, red, Fig. 6B; Particle Image Velocimetry (PIV), Supp. Fig. 5B) translates to compressive stress on the progerin cell. Conversely, the traction force pointing toward the progerin cell (blue, Fig. 6B) represents tensile stress. The temporal profile of the average radial traction force (Methods, Fig. 6C, G) showed persistent positive radial traction force around progerin-expressing cells, indicating that these cells experienced persistent compressive force prior to elimination. This positive traction force diminished at the onset of cell elimination, consistent with the disruption of cell-substrate adhesion upon apoptosis. The persistent positive radial traction force prior to elimination was in stark contrast to the control condition, where the traction forces were on average weaker (Fig. 6D-F, Supp. Video 6, PIV, Supp. Fig. 5D). This was further supported by the average behavior of multiple samples. Cells surrounding progerin-expressing cells consistently showed positive radial traction force (Fig. 6H). In contrast, cells surrounding control cells showed inconsistent radial traction force, with some samples showing positive and others negative, resulting in an average around zero (Fig. 6I). Collectively, quantitative assessment of traction force around progerin-expressing cells showed that mechanical compression by neighboring cells precedes the elimination of progerin-induced senescent cells in co-culture, supporting the idea of mechanical cell competition.

**Fig. 6.**
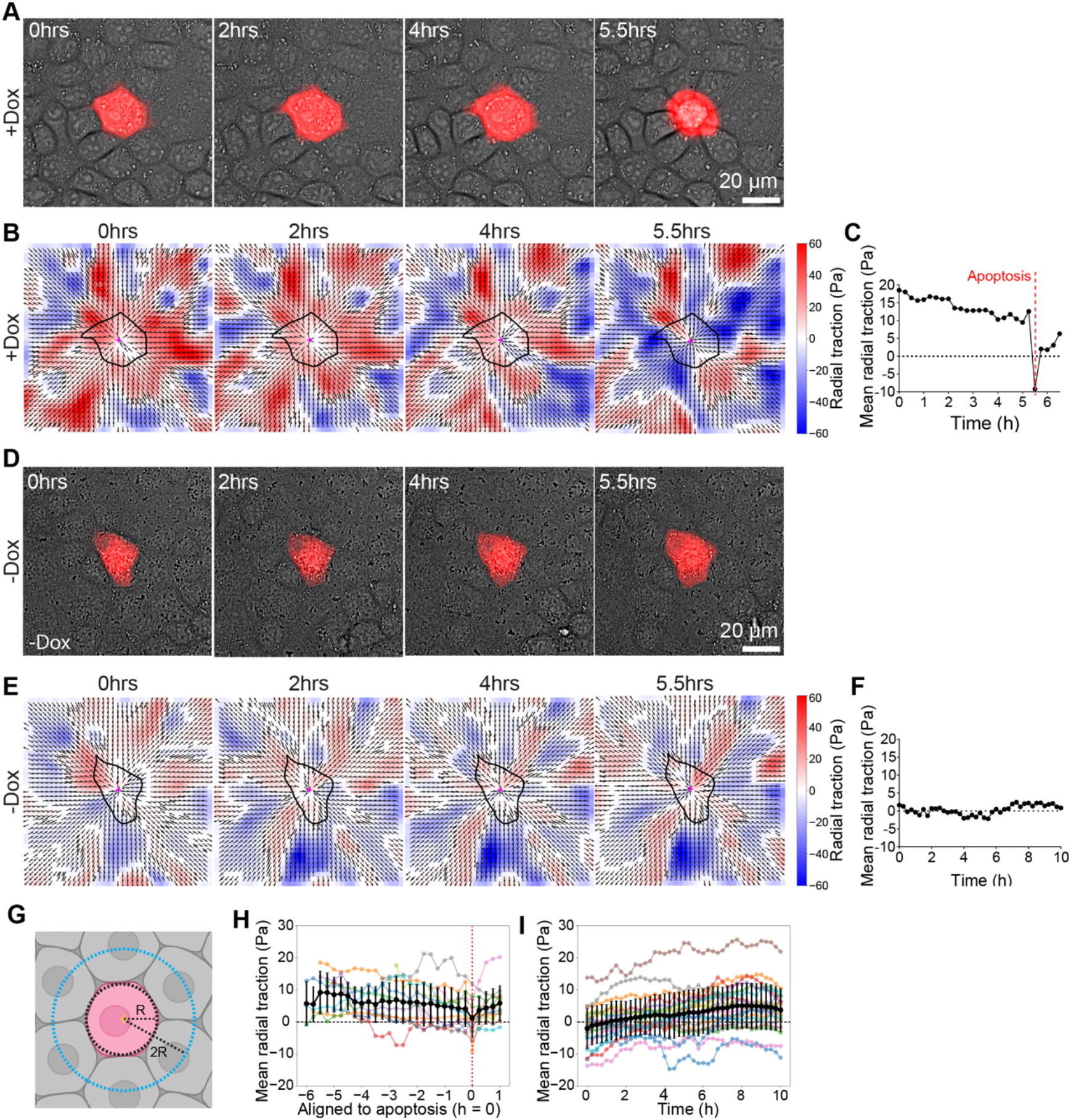
Mechanical compression by neighboring cells precedes apoptosis of progerin-expressing cells. (A) Representative time-lapse images of mCherry-Progerin cells (red) co-cultured with normal cells with doxycycline treatment. (B) Representative radial traction maps of co-culture tissue with doxycycline treatment at indicated time points. The progerin cell is outlined. Red and blue indicate radial traction force pointing away from (compressive) and toward (tensile) the progerin cell, respectively. (C) Time course of mean radial traction in the neighboring region (R–2R) for the sample shown in (B). (D) Representative time-lapse images of mCherry-Progerin cells (red) co-cultured with normal cells without doxycycline treatment. (E) Representative radial traction maps of co-culture tissue without doxycycline treatment. (F) Time course of mean radial traction for the sample shown in (E). (G) Schematic of annular regions used for radial traction analysis. R is the equivalent circular radius of the progerin cell. (H) Mean radial traction over time aligned to the time of apoptosis (t = 0) from multiple co-culture samples with doxycycline treatment. Individual samples are shown in color; the average is shown in black. (I) Mean radial traction over time from multiple co-culture samples without doxycycline treatment. Data are presented as mean ± SD. In (H), (independent repeats, n = 5, N = 13); In (I), (independent repeats, n = 3, N = 21).

### p16-induced senescent cells are resistant to apoptosis in co-culture tissue

To address whether the observed elimination of senescent cells is specific to progerin-induced senescence, we sought to induce cellular senescence through a pathway independent of DNA-damage and p53 pathways^30^, which is the pathway through which progerin induces cellular senescence. To this end, we established a cell line overexpressing p16 using a doxycycline-inducible system (mNG-T2A-p16, hereafter referred to as mNG-p16, Fig. S6A). mNG-p16 cells exhibited characteristics of cellular senescence upon three days of doxycycline treatment (Fig. S6B-E). Live imaging of mNG-p16 cells co-cultured with normal cells showed that most mNG-p16 cells persisted in the tissue for at least two days (Fig. 7A-B, Supp. Video 7), unlike mNG-Progerin cells under the same co-culture condition. To understand the difference between these two conditions, we evaluated the levels of JNK and p-p38-MAPK by immunostaining. The levels of JNK and p-p38-MAPK in mNG-p16 cells were higher than in neighboring normal cells, similar to our observations in mNG-Progerin cells, yet lower than in p16 cells cultured alone (Fig. 7C-H). Notably, the level of p21 in mNG-p16 cells under co-culture was higher than in neighboring normal cells (Fig. 7I-K), unlike the reduction of p21 observed in mNG-Progerin cells under co-culture conditions. The levels of JNK, p-p38-MAPK, and p21 without doxycycline treatment were similar to the control condition (Fig. S7A-I). These findings suggest that p16-induced senescent cells are resistant to apoptosis in the presence of high levels of p21, which aligns with previous studies showing that senescent cells, including p16-positive cells, accumulate in tissues during aging^31^. This contrast in cellular fate between progerin-induced and p16-induced senescent cells highlights that the non-cell-autonomous elimination of senescent cells is context-dependent.

**Fig. 7.**
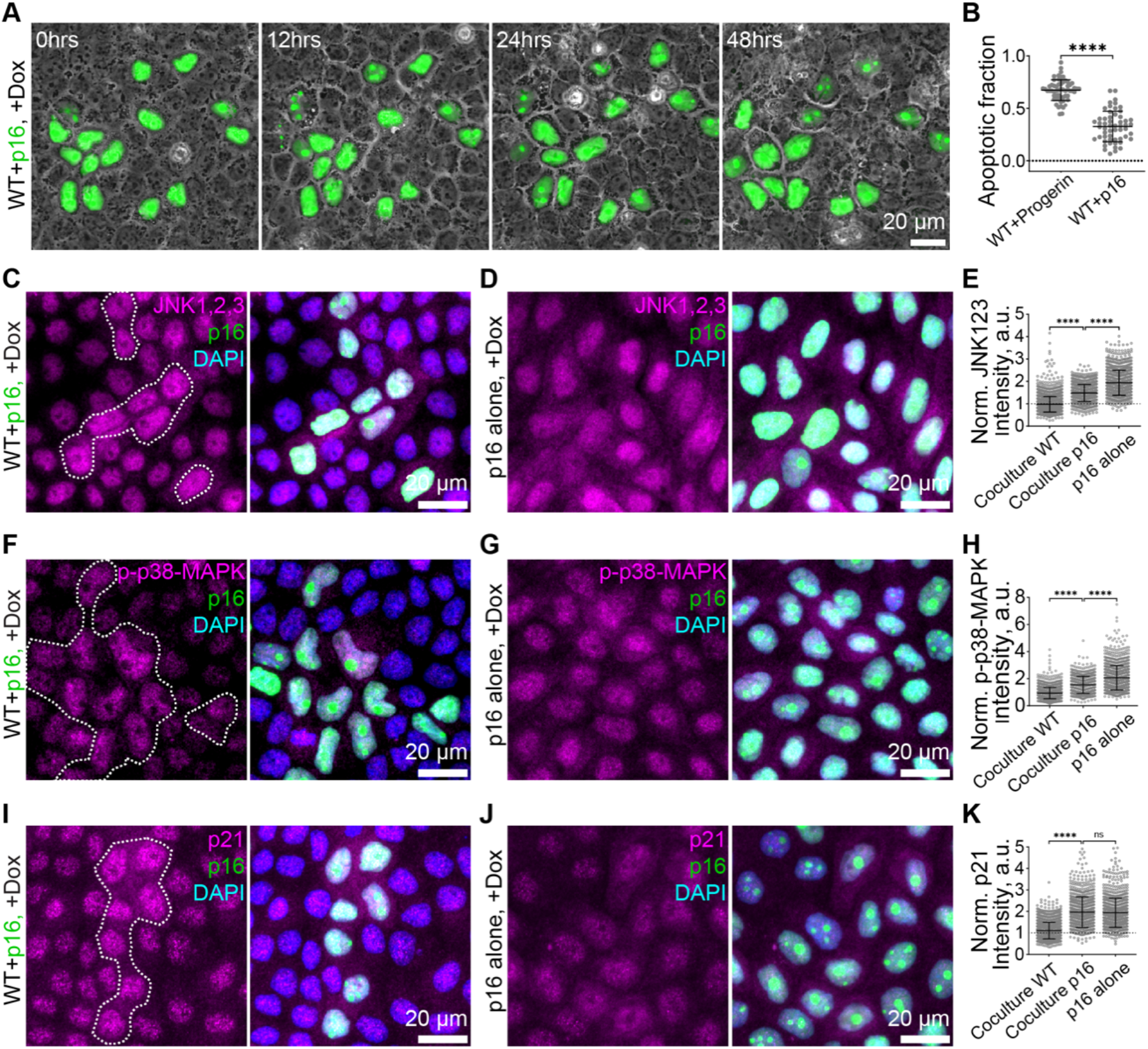
p16-induced senescent cells are resistant to apoptosis in co-culture tissue. (A) Time-lapse images of mNG-p16 cells (green) co-cultured with normal cells after doxycycline treatment. (B) Apoptotic fraction of mNG-Progerin cells (shown in Figure1D) and mNG-p16 cells in co-culture with normal cells. (C) Representative immunofluorescence images of JNK1,2,3 (magenta) in mNG-p16 co-culture tissue with doxycycline treatment. p16 cells are indicated by dashed outlines. Nuclei were stained with Hoechst 33342 (blue). (D) Representative immunofluorescence images of JNK1,2,3 in p16 cells cultured alone with doxycycline treatment. (E) Quantification of normalized JNK1,2,3 fluorescence intensity in neighboring normal cells and p16 cells in co-culture, and p16 cells cultured alone. (F) Representative immunofluorescence images of p-p38-MAPK (magenta) in mNG-p16 co-culture tissue with doxycycline treatment. (G) Representative immunofluorescence images of p-p38-MAPK in p16 cells cultured alone with doxycycline treatment. (H) Quantification of normalized p-p38-MAPK fluorescence intensity in neighboring normal cells and p16 cells in co-culture, and p16 cells cultured alone. (I) Representative immunofluorescence images of p21 (magenta) in mNG-p16 co-culture tissue with doxycycline treatment. (J) Representative immunofluorescence images of p21 in p16 cells cultured alone with doxycycline treatment. (K) Quantification of normalized p21 fluorescence intensity in neighboring normal cells and p16 cells in co-culture, and p16 cells cultured alone. Data are presented as mean ± SD. An unpaired two-tailed t-test is used in (B), (independent repeats, n = 3, N = 60 and 55). Kruskal-Walli’s test and Dunn’s multiple comparisons test are used in (E), (independent repeats, n = 3 (for coculture) and n = 4 (for p16 alone), N = 2099, 1133, and 992). Kruskal-Walli’s test and Dunn’s multiple comparisons test are used in (H), (independent repeats, n = 3 (for coculture) and n = 4 (for p16 alone), N = 1513, 676, and 1207). Kruskal-Walli’s test and Dunn’s multiple comparisons test were used in (K), (independent repeats, n = 3 (for coculture) and n = 4 (for p16 alone), N = 1490, 842, and 1023). ns, not significant; ^**^*p*<0.0001.

## Discussion

In this study, we found that progerin-induced senescent cells undergo apoptosis when surrounded by normal cells in co-culture. This elimination requires direct cell-cell contact and is mediated by the JNK and p38-MAPK pathways, leading to p53 upregulation and p21 downregulation in progerin-expressing cells. Traction force microscopy revealed that neighboring normal cells exert persistent mechanical compression on progerin-expressing cells prior to their elimination, consistent with mechanical cell competition. In contrast, p16-induced senescent cells resist elimination under the co-culture conditions, highlighting that the non-cell-autonomous elimination of senescent cells is context-dependent.

A key finding of this study is the reduction of p21 in progerin-expressing cells under co-culture conditions. p21 is typically elevated in senescent cells, where it functions as a key mediator of cell cycle arrest and confers resistance to apoptosis^29,32^. However, our data showed that p21 levels in progerin-expressing cells decreased specifically in the co-culture context, not when progerin-expressing cells were cultured alone. This reduction was partially rescued by JNK and p38 inhibitors, suggesting that the JNK and p38-MAPK pathways activated by neighboring normal cells contribute to the downregulation of p21. The resulting decrease in p21 may shift the balance from cell cycle arrest and apoptosis resistance toward apoptosis, effectively overriding the typical apoptosis-resistant phenotype of senescent cells.

One notable question arising from our findings is what determines the contrasting fates of progerin-induced and p16-induced senescent cells in co-culture. Progerin overexpression induces cellular senescence through the DNA damage response and p53 pathway activation^5^, whereas p16 overexpression induces cell cycle arrest through the p16^INK4A^-Rb axis, independent of the DNA damage response^3^. We speculate that this fundamental difference in senescence pathways underlies the distinct outcomes^33^. In progerin-expressing cells, the activation of JNK and p38-MAPK pathways by neighboring cells leads to p21 downregulation and subsequent apoptosis. In contrast, p16-induced senescent cells maintain high p21 levels under co-culture conditions despite elevated JNK and p38-MAPK, suggesting that the p16^INK4A^ pathway sustains p21 expression through a mechanism^34,35^ that is resistant to JNK and p38-mediated downregulation. Previous studies have demonstrated that cells with DNA damage can be eliminated from tissues^36,37^, which is consistent with the selective vulnerability of progerin-induced senescent cells observed here.

Although our data demonstrated that long-range secreted factors are unlikely to be the primary driver of progerin-induced cell elimination, we cannot rule out a modulatory role for SASP in this process. Previous studies have shown that SASP profiles exhibit high heterogeneity between p53/p21-induced and p16-induced senescent cells^33,38^, and that progerin expression triggers a p53-mediated SASP^39^. It is possible that differences in SASP composition between progerin-induced and p16-induced senescent cells influence how neighboring normal cells respond, potentially modulating the mechanical forces exerted on senescent cells. Further investigation of how SASP profiles shape the mechanical interactions between senescent and normal cells will be an important direction for future studies.

Our study has several limitations. Our findings are based on an *in vitro* co-culture system, and whether the mechanical cell competition observed here operates in tissues *in vivo* remains to be determined. In living tissues, senescent cells exist within a complex three-dimensional microenvironment with extracellular matrix^40^, immune cells^41^, and diverse cell types^42^, all of which may influence the elimination process. Additionally, while we identified JNK, p38-MAPK, p53, and p21 as key factors, the precise molecular mechanisms by which neighboring cells sense and mechanically compress progerin-expressing cells require further investigation.

Together, our findings reveal a non-cell-autonomous mechanism for the clearance of senescent cells through mechanical cell competition. This mechanism complements the known immune-mediated clearance of senescent cells and provides additional insight into how tissues maintain homeostasis during aging. Understanding these intercellular interactions may offer new avenues for therapeutic strategies aimed at selectively removing senescent cells in age-related diseases.

## References

1. López-Otín, C., Blasco, M. A., Partridge, L., Serrano, M. & Kroemer, G. Hallmarks of aging: An expanding universe. Cell 186, 243–278 (2023).

2. Hu, L. et al. Why Senescent Cells Are Resistant to Apoptosis: An Insight for Senolytic Development. Front Cell Dev Biol 10, 822816 (2022).

3. Ajoolabady, A. et al. Hallmarks of cellular senescence: biology, mechanisms, regulations. Exp Mol Med 57, 1482–1491 (2025).

4. Gonzalo, S., Kreienkamp, R. & Askjaer, P. Hutchinson-Gilford Progeria Syndrome: a premature aging disease caused by LMNA gene mutations. Ageing Res Rev 33, 18–29 (2017).

5. Cadiñanos, J., Varela, I., López-Otín, C. & Freije, J. M. P. From immature lamin to premature aging: molecular pathways and therapeutic opportunities. Cell Cycle 4, 1732–1735 (2005).

6. Wang, B., Han, J., Elisseeff, J. H. & Demaria, M. The senescence-associated secretory phenotype and its physiological and pathological implications. Nat Rev Mol Cell Biol 25, 958–978 (2024).

7. Chou, L.-Y.Ho, C.-T. & Hung, S.-C. Paracrine Senescence of Mesenchymal Stromal Cells Involves Inflammatory Cytokines and the NF-κB Pathway. Cells 11, 3324 (2022).

8. Krishnan, S., Paul, P. K. & Rodriguez, T. A. Cell competition and the regulation of protein homeostasis. Current Opinion in Cell Biology 87, 102323 (2024).

9. Baker, N. E. Emerging mechanisms of cell competition. Nat Rev Genet 21, 683–697 (2020).

10. Baker, N. E. Mechanisms of cell competition emerging from Drosophila studies. Curr Opin Cell Biol 48, 40–46 (2017).

11. Kajita, M. et al. Filamin acts as a key regulator in epithelial defence against transformed cells. Nat Commun 5, 4428 (2014).

12. Hogan, C. et al. Characterization of the interface between normal and transformed epithelial cells. Nat Cell Biol 11, 460–467 (2009).

13. Kon, S. et al. Cell competition with normal epithelial cells promotes apical extrusion of transformed cells through metabolic changes. Nat Cell Biol 19, 530–541 (2017).

14. Kohashi, K. et al. Sequential oncogenic mutations influence cell competition. Current Biology 31, 3984–3995.e5 (2021).

15. Yamamoto, M., Ohsawa, S., Kunimasa, K. & Igaki, T. The ligand Sas and its receptor PTP10D drive tumour-suppressive cell competition. Nature 542, 246–250 (2017).

16. Levayer, R. Cell Competition: How to Take Over the Space Left by Your Neighbours. Current Biology 28, R741–R744 (2018).

17. Soares, C. C., Rizzo, A., Maresma, M. F. & Meier, P. Autocrine glutamate signaling drives cell competition in Drosophila. Developmental Cell 59, 2974–2989.e5 (2024).

18. Alpar, L., Bergantiños, C. & Johnston, L. A. Spatially Restricted Regulation of Spätzle/Toll Signaling during Cell Competition. Dev Cell 46, 706–719.e5 (2018).

19. Schoenit, A. et al. Force transmission is a master regulator of mechanical cell competition. Nat. Mater. 24, 966–976 (2025).

20. Wagstaff, L. et al. Mechanical cell competition kills cells via induction of lethal p53 levels. Nat Commun 7, 11373 (2016).

21. Bertheloot, D., Latz, E. & Franklin, B. S. Necroptosis, pyroptosis and apoptosis: an intricate game of cell death. Cell Mol Immunol 18, 1106–1121 (2021).

22. Chojnowski, A. et al. Heterochromatin loss as a determinant of progerin-induced DNA damage in Hutchinson-Gilford Progeria. Aging Cell 19, e13108 (2020).

23. Saw, T. B. et al. Transepithelial potential difference governs epithelial homeostasis by electromechanics. Nat. Phys. 18, 1122–1128 (2022).

24. Senoo-Matsuda, N. & Johnston, L. A. Soluble factors mediate competitive and cooperative interactions between cells expressing different levels of Drosophila Myc. Proceedings of the National Academy of Sciences 104, 18543–18548 (2007).

25. Guo, Y.-J. et al. ERK/MAPK signalling pathway and tumorigenesis. Exp Ther Med 19, 1997–2007 (2020).

26. Pua, L. J. W. et al. Functional Roles of JNK and p38 MAPK Signaling in Nasopharyngeal Carcinoma. International Journal of Molecular Sciences 23, 1108 (2022).

27. Canovas, B. & Nebreda, A. R. Diversity and versatility of p38 kinase signalling in health and disease. Nat Rev Mol Cell Biol 22, 346–366 (2021).

28. Xu, R. & Hu, J. The role of JNK in prostate cancer progression and therapeutic strategies. Biomedicine & Pharmacotherapy 121, 109679 (2020).

29. Mirzayans, R., Andrais, B., Hansen, G. & Murray, D. Role of p16INK4A in Replicative Senescence and DNA Damage-Induced Premature Senescence in p53-Deficient Human Cells. Biochemistry research international 2012, 951574 (2012).

30. Prieur, A., Besnard, E., Babled, A. & Lemaitre, J.-M. p53 and p16INK4A independent induction of senescence by chromatin-dependent alteration of S-phase progression. Nat Commun 2, 473 (2011).

31. Lyons, C. E. et al. Chronic social stress induces p16-mediated senescent cell accumulation in mice. Nat Aging 5, 48–64 (2025).

32. Yan, J. et al. The role of p21 in cellular senescence and aging-related diseases. Mol Cells 47, 100113 (2024).

33. Saul, D. et al. Distinct senotypes in p16-and p21-positive cells across human and mouse aging tissues. EMBO J 44, 7295–7325 (2025).

34. Al-Mohanna, M. A., Al-Khalaf, H. H., Al-Yousef, N. & Aboussekhra, A. The p16 INK4a tumor suppressor controls p21 WAF1 induction in response to ultraviolet light. Nucleic Acids Res 35, 223–233 (2007).

35. Al-Khalaf, H. H. & Aboussekhra, A. p16INK4A Positively Regulates p21WAF1 Expression by suppressing AUF1-dependent mRNA decay. PLOS ONE 8, e70133 (2013).

36. Kato, T. et al. Dynamic stem cell selection safeguards the genomic integrity of the epidermis. Dev Cell 56, 3309–3320.e5 (2021).

37. Kozyrska, K. et al. p53 directs leader cell behavior, migration, and clearance during epithelial repair. Science 375, eabl8876 (2022).

38. Sturmlechner, I. et al. p21 produces a bioactive secretome that places stressed cells under immunosurveillance. Science 374, eabb3420 (2021).

39. Manakanatas, C. et al. Endothelial and systemic upregulation of miR-34a-5p fine-tunes senescence in progeria. Aging (Albany NY) 14, 195–224 (2022).

40. Mavrogonatou, E., Pratsinis, H., Papadopoulou, A., Karamanos, N. K. & Kletsas, D. Extracellular matrix alterations in senescent cells and their significance in tissue homeostasis. Matrix Biol 75–76, 27–42 (2019).

41. Majewska, J. & Krizhanovsky, V. Immune surveillance of senescent cells in aging and disease. Nat Aging 5, 1415–1424 (2025).

42. Gurkar, A. U. et al. Spatial mapping of cellular senescence: emerging challenges and opportunities. Nat Aging 3, 776–790 (2023).

43. Foo, M. X. R. et al. Genetic and pharmacological modulation of lamin A farnesylation determines its function and turnover. Aging Cell 23, e14105 (2024).

44. Kawaue, T. et al. Inhomogeneous mechanotransduction defines the spatial pattern of apoptosis-induced compensatory proliferation. Dev Cell 58, 267–277.e5 (2023).

45. Park, J. H. et al. Materials and extracellular matrix rigidity highlighted in tissue damages and diseases: Implication for biomaterials design and therapeutic targets. Bioact Mater 20, 381–403 (2023).

46. Teo, J. L., Lim, C. T., Yap, A. S. & Saw, T. B. A Biologist’s Guide to Traction Force Microscopy Using Polydimethylsiloxane Substrate for Two-Dimensional Cell Cultures. STAR Protoc 1, 100098 (2020).

47. Mori, Y. et al. Extracellular ATP facilitates cell extrusion from epithelial layers mediated by cell competition or apoptosis. Current Biology 32, 2144–2159.e5 (2022).

48. Schneider, C. A., Rasband, W. S. & Eliceiri, K. W. NIH Image to ImageJ: 25 years of image analysis. Nat Methods 9, 671–675 (2012).

